# Deep Active Learning for Robust Biomedical Segmentation

**DOI:** 10.1101/2023.03.28.534521

**Authors:** Mustafa Arikan, Ferenc Sallo, Andrea Montesel, Hend Ahmed, Ahmed Hagag, Marius Book, Henrik Faatz, Maria Cicinelli, Sepehr Meshkinfamfard, Sevim Ongun, Adam Dubis, Watjana Lilaonitkul

## Abstract

Deep learning for medical applications faces many unique challenges. A major challenge is the large amount of labelled data for training, while working in a relatively data scarce environment. Active learning can be used to overcome the vast data need challenge. A second challenged faced is poor performance outside of a experimental setting, contrary to the high requirement for safety and robustness. In this paper, we present a novel framework for estimating uncertainty metrics and incorporating a similarity measure to improve active learning strategies. To showcase effectiveness, a medical image segmentation task was used as an exemplar. In addition to faster learning, robustness was also addressed through adversarial perturbations. Using epistemic uncertainty and our framework, we can cut number of annotations needed by 39% and by 54% using epistemic uncertainty and a similarity metric.

## 1 Introduction

Deep learning (DL) for medical applications has shown great promise in experimental settings, but results have not generalized broadly, resulting in poor performance on real-world validation tests[2, 12, 4]. One key attributing factor is the difficulty in obtaining labels for medical data in large enough quantities for training. While data availability is rapidly increasing, aggregation remains difficult due to legality and data sensitivity. Labelled training data can be difficult to obtain due to the time and expertise required for production. One solution to overcome the latter challenge is via ‘active learning’[3]. Active learning entails a system learning from data that automatically prioritises what additional data it needs for experts to label, in order to increase learning efficiencies while minimizing labelling efforts. With each active learning iteration, an acquisition function with a selection criteria such as model uncertainty is used to rank and select data samples that are prioritized for labelling and addition to the training set. For active learning with DL models, [6] used the dropout method as a Bayesian approximation of the epistemic uncertainty (or model’s uncertainty)[5, 9, 10] as the selection criteria to achieve state-of-the-art (SOTA) performance in active learning on classification tasks.

However, using model uncertainty as a selection criterion by itself can be limiting, especially when the selected data samples map to a small subset of similar images. This can induce a lower diversity over the training set and potentially reduce the trained model’s robustness. A model is more robust when its accuracy is not very sensitive to adversarial data samples that contain perturbations, such as variations in noisiness or changes in gamma corrections to medical images acquired from different machine-makes. Previous studies have examined the trade-off between accuracy and robustness in classification[13, 17, 16]. To date, most studies on DL in medicine and healthcare have focused mostly on accuracy as the performance metric. We argue that the measure of model robustness is an important additional performance indicator to attain some limits on the safety-of-use of DL models applied to medicine where prediction mistakes can be costly to human health.

In this study, we propose an active learning framework that aims to concurrently optimize data-efficiency, performance accuracy and model robustness. To ensure sample selection diversity during active learning, we propose the use of image similarity as a metric to cluster the unlabeled data and then prioritise the data within each cluster based on epistemic uncertainty. We quantified the incremental gains in robustness and accuracy from our active learning framework on the task of retinal layer segmentation in optical coherence tomography (OCT) images against two baseline learning frameworks: (i) learning with random sample-selection and (ii) SOTA active learning using epistemic uncertainty[7, 8]. We demonstrated that our method outperforms both baselines in data-efficiency and performance accuracy (see figure 1) and in robustness (see figure 2). In addition, in a transferred learning application, using trained models from our method to fine-tune on images from a different and unseen disease gave higher performance than using trained models from random sampling at all training set sizes (see figure 3).

**Figure 1.**
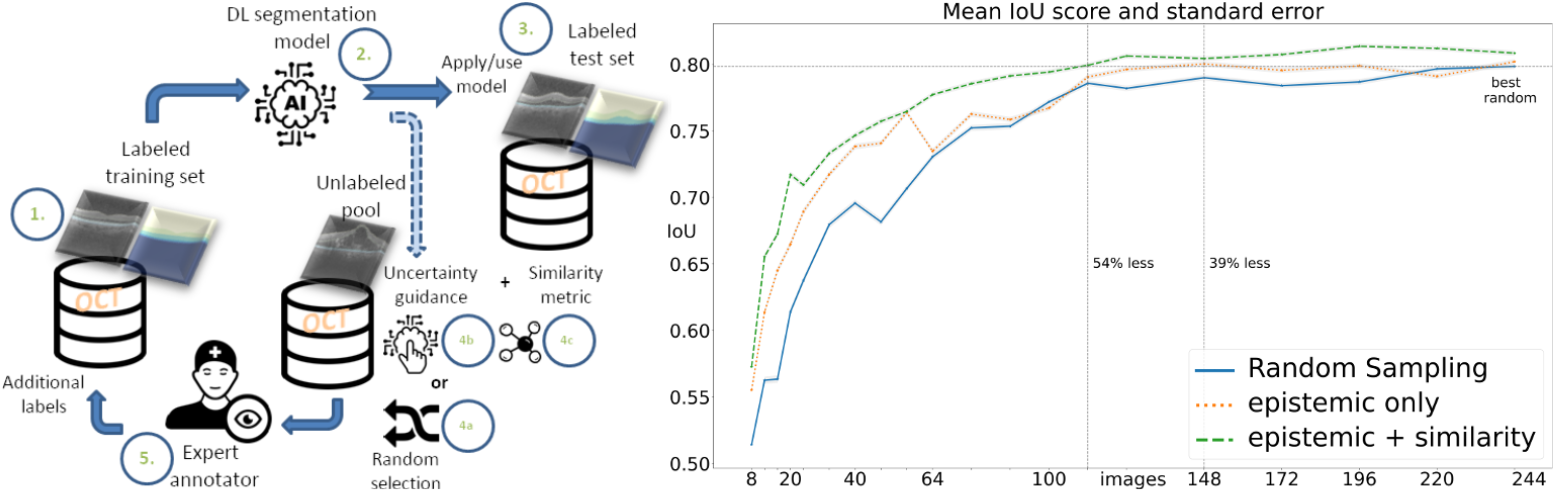
Our deep active learning approach using similarity and model uncertainty to improve robustness and predictive performance while minimizing labelling efforts of the training data. (Left) shows an illustration of the process of adding samples to the training set.(Right) shows the intersection-over-union (IoU) results averaged from 10 experimental repeats from 3 experiments: (i) random sample selection (blue), (ii) SOTA method using epistemic uncertainty only (orange) [7], and (iii) our proposed method of combining similarity and epistemic uncertainty (green).

**Figure 2.**
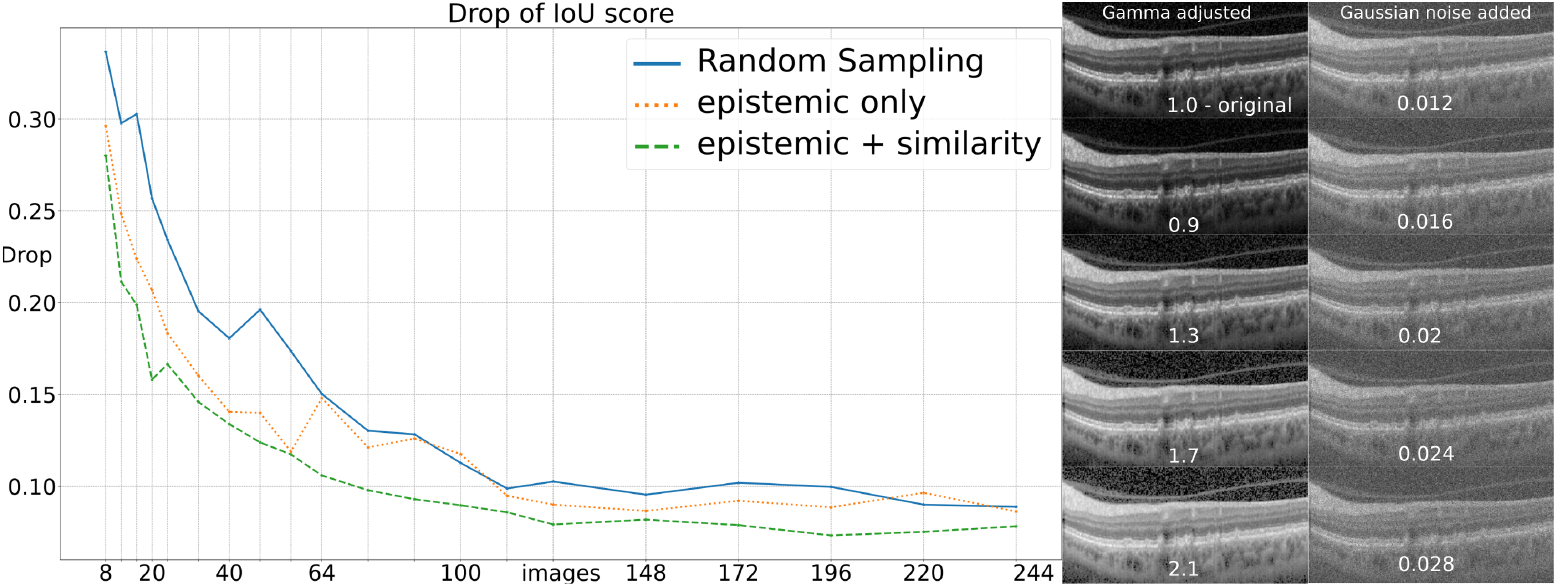
Right: perturbation of a sample test image with gamma adjustment (first column), from top to bottom: 1.0, 0.7, 0.9, 1.1, 1.3, 1.5, 1.7, 1.9, 2.1, 2.3 and 2.5 and Gaussian noise (second row) with variance: 0.01, 0.012, 0.014, 0.016, 0.018, 0.02, 0.022, 0.024, 0.026 and 0.028. Left: Drop of IoU accuracy compared with test set and extended test set containing perturbed examples using random selection, epistemic uncertainty only, and combined with similarity.

**Figure 3.**
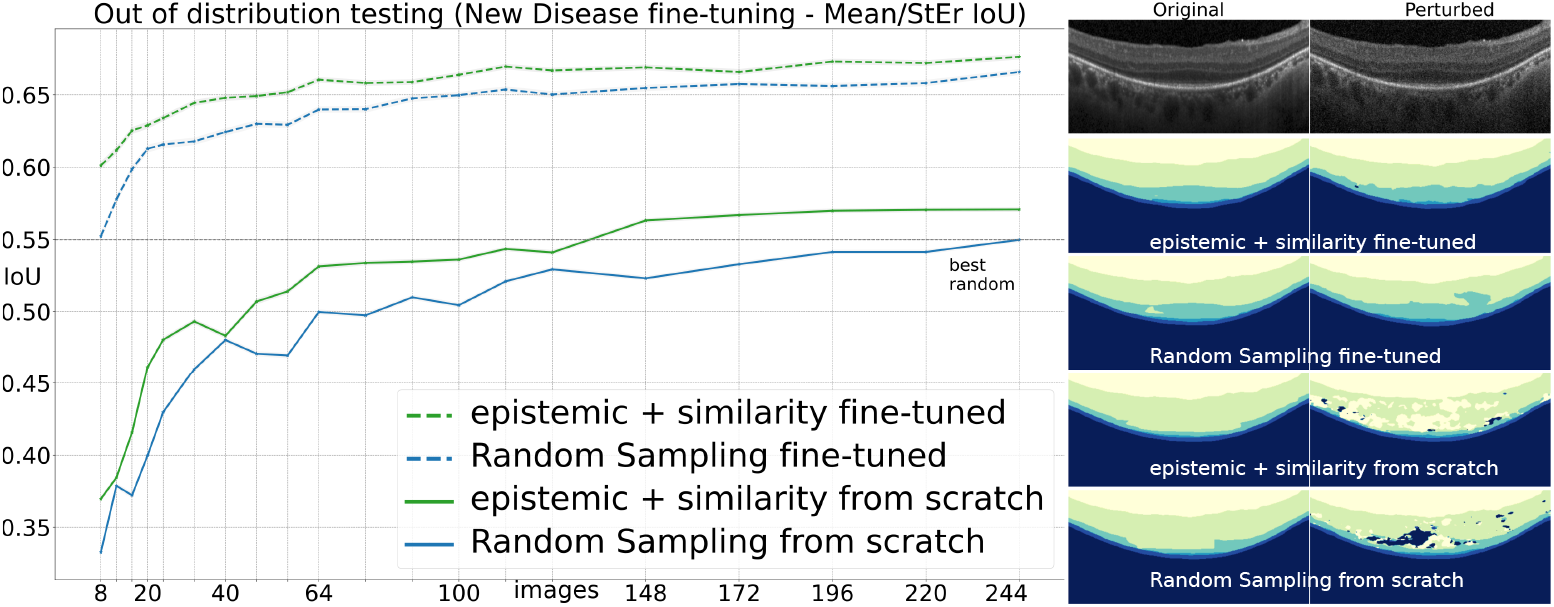
Fine-tuning the segmentation model with different learning strategies on a new disease dataset with reported test-time performance on adversarial samples. (Left) shows IoU performance with training from scratch (1 - blue/bottom), fine-tuning on the random selection from the original data set with random selection (2 - blue/top), training with epistemic and similarity from scratch (3 - green/bottom) and fine-tuning on active learning with epistemic uncertainty and similarity (4 - green/top). (Right): one qualitative example predicted with 1,2,3 and 4 at 12th iteration with 100 images added for an original image and a perturbed scan.

## 2 Methodology

### Data sets

The first dataset[11] consists of AMD (Age-related Macular Degeneration), DME (Diabetic Macular Edema), and healthy OCT scans. We split our pool of OCT volumes representing different patients (60 volumes and 1668 scans in total with equal distribution across classes) into 20% training, 20% validation, and 60% testing. The second private data set consists of 2112 scans from 44 patients with Usher syndrome with the same splits as the main data set. The images were segmented manually for the presence and location of key laminar boundaries by multiple experts for quality control of the labels.

### Experiments

We carried out two experiments using three different acquisition functions for iterative learning on the task of on the task of retinal layer segmentation with U-Net and EfficientNet as the network backbone[15]. The three acquisition functions are (i) random selection (no active learning), (ii) a SOTA active learning method using epistemic uncertainty only[7, 8], and (iii) active learning with similarity and epistemic uncertainty (ours). We reported the average results from 10 repeats for each experiment. In the first experiment, we quantified the incremental gains in data-efficiency and prediction performance measured by the intersection-over-union (IoU) metric (see figure 1) at different training sample sizes from data set 1. The second experiment investigated the robustness of the trained models with adversarial test samples, which were generated by adding random perturbations using a range of gamma adjustments and Gaussian noise (see figure 2). For our proposed active learning setup, we combined a similarity metric and epistemic uncertainty to compose an acquisition function. The similarity metric (SSIM)[14] was used to calculate a pairwise distance matrix between all images. The distance matrix was used to create clusters of similar images using k-means[1]. To perform active learning and query new examples within each cluster, we employed Monte Carlo dropout[5] to estimate the epistemic or model uncertainty associated with each unlabelled image and then selected samples with the highest values. The numbers of samples or images added during the iterations ranged from 8 to 244. See the left pane of figure 1 for the iterative learning setup.

### Application to transferred learning

To investigate how well our proposed method can transfer knowledge learned onto a new dataset, we compared the performance between our method with the random sampling baseline where the models were first pre-trained using data set 1 and then fine-tuned with images from data set 2, which contained images from a different disease.

## 3 Results

Results show that our method outperforms both baselines in data-efficiency and performance accuracy (see figure 1). At the best performance level achieved by random sampling, our method reduces the required annotated data by 54% whereas the SOTA baseline method reduced the required annotated data by 39%. Our method also achieves a higher prediction performance at all sizes of training data against both baselines. In terms of robustness, the performance of models trained from our method were less affected by adversarial test examples at all training set sizes than models from both baseline models (see figure 2). Transferred learning to images from a different disease by fine-tune training (see dotted lines in figure 3) gave higher performance than when training the models from scratch (see solid lines in figure 3) for both our proposed method and the random sampling baseline. In the fine-tune learning experiments, our method outperformed the random sampling baseline consistently across all training sample sizes.

## 4 Discussion

Medical image annotation is a time-intensive task which can limit the ability to train DL models for medical applications. Our active learning framework outlines a data labelling pipeline that ensures diversity of sample selection while minimizing model uncertainty. This simple strategy significantly reduced labelling efforts to achieve a desired prediction performance. More importantly, by ensuring sample diversity for active learning, we show that model robustness is also significantly improved.

Increased robustness is an important aspect to optimize especially in medical-related tasks that can directly impact human health. Lastly, we also demonstrated training efficiencies and performance gain when using models trained from our method to fine-tune on a new dataset from a different disease. In this respect, the combination of similarity with uncertainty for training data selection can help reduce the burden of manual annotation while improving the performance and robustness by ensuring a higher level of diversity in the selected data for training.

## 5 Conclusion and Future Work

We have introduced a new framework for uncertainty-based medical image segmentation for Deep Bayesian Active Learning where diversity of samples is enhanced using a similarity metric. Our method can significantly reduce annotation effort while outperforming the baseline methods on both prediction performance and model robustness measures. In future work, we would like to understand the applicability of our approach to other adversarial and out-of-distribution examples and different tasks such as classification and detection. We also aim to extend the framework to metrics beyond similarity and epistemic uncertainty as well as investigate data-driven ways to better combine the metrics for active learning.

## 6 Potential Negative Societal Impacts

In medical imaging and in medicine deep learning models require measures for safety, robustness, and interpretability. Deep learning models are considered black-box models where the output often cannot be fully explained, which is unhelpful in safety-critical domains such as medical imaging. Furthermore, the comparison between medical experts and AI systems can be misguiding. Deep learning methods need to fit into a framework where domain experts can process and consider the sources of uncertainty interactively to understand and mitigate risks.

## Acknowledgements

The authors would also like to acknowledge the use of the Joint Academic Data science Endeavour (JADE) Tier 2 computing facility funded by the Engineering and Physical Sciences Research Council (EPSRC), UK.

